# A structural transition ensures robust formation of skeletal muscle

**DOI:** 10.1101/2025.09.22.677369

**Authors:** Mario A. Mendieta-Serrano, Yiqi Hou, Sophie Theis, Thomas E. Hall, Shannon E. Taylor, Berta Verd, Robert G. Parton, Timothy E. Saunders

## Abstract

During organ development cells undergo significant morphological and positional changes. Yet, by the end of organogenesis, internal organ structure is typically robustly defined, with cells tightly packed. It remains an open question as to how the three-dimensional (3D) internal structure of an organ emerges reliably, particularly when there are multiple cell types interacting and dynamic boundary constraints. Here, we utilise quantitative live imaging and 3D morphological measures of the developing zebrafish myotome to unravel how early muscle organisation emerges. Contrary to the textbook view of muscle fibres as cylindrical, myocytes undergo an ordered chiral twist, the direction and magnitude of which depends on their position within the myotome. Further, cells skew and rearrange, seemingly to facilitate close packing of neighbouring muscle fibres. Cell movement undergoes a rapid decline in speed once the cells span the myotome segment. We find that cell packing is altered in mutants that disrupt cell fate or cell fusion, even though the final muscle segments remain largely confluent. Biophysical perturbation reveals that the cells are mechanically plastic, able to adjust to changes in the local cellular environment and boundary constraints. Taking these results together, we propose that the early myotome undergoes a structural transition, from a fluid-like state into a frozen state, resembling glass-like behaviour. Cellular plasticity in response to varying boundary constraints may be a general mechanism for ensuring robust organ morphogenesis in dense 3D tissues.

## Introduction

Internal organs have multiple cell types and complex spatial arrangement. Due to this complexity, it remains an open challenge in developmental biology to decipher the mechanisms by which internal organs robustly form in most systems. The formation of external organs, such as skin morphogenesis or hair placodes, has revealed how long-range biomechanical processes can shape organs (1–3). Plant systems have offered key insights into the interplay between cell behaviours and organ shape (4, 5); the slower dynamics and (typically) clearer cells making quantitative analysis more accessible (6). Internal organ morphogenesis in animal systems remains comparatively poorly understood, especially at sufficient spatiotemporal scales to explore how multiple processes interact.

Cells have to adapt to both local (*e.g*., neighbouring cells) and global (*e.g*., boundaries) constraints during organogenesis (7–10). These constraints can significantly affect cell shape, as recently demonstrated in a range of organs and tissues (11–15). Further, regulation of cell motility within tissues can result in abrupt changes in material properties; cells can transition between fluid-like (able to readily exchange neighbours) to solid-like behaviour (rigid) (10, 16). This behaviour has drawn comparisons with phase transitions, with both jamming (17, 18) and percolation (19) transitions described during early embryonic development. Glass-like transitions have also been observed in biological systems (20); cells undergo a rapid slow down in motility and fix into a rigid state where the final configuration depends on the particular morphodynamics in each case. It has been proposed that such material transitions potentially play a role in ensuring robust organ formation and wound repair (21, 22), but this has yet to be demonstrated in complex 3D organs *in vivo*.

Analysis of cell morphodynamics within confluent tissues has largely been in systems that can be approximated as 2D or in systems with a single cell type and relatively simple boundary constraints (17, 19, 23–25). Formation of internal 3D structure has been studied in tubular organs such as the lung and kidneys. However, such work has mainly focused on how the branching network structures emerge (26–28), rather than the processes driving shape formation at a cell scale (29). Single cell sequencing methods have revealed that cell fates are precisely determined in space and time during organ development (30–32), including in humans (33–35). These approaches have been combined with high resolution imaging (36) to build cell-scale atlases of organ development (37, 38). However, these methods are still static, at least on the typical time scale of cell morphogenesis. For example, skeletal muscle fibres can transition from approximately spherical to highly elongated in less than one hour (39, 40).

Here, we investigate how cells shape and pack in 3D within a dense tissue with multiple cell types and spatiotemporally varying boundary constraints. We focus on the developing skeletal muscle in zebrafish. The muscle is formed of slow and fast muscle fibres (Fig. 1A) (41–43), derived from myocyte progenitors. The skeletal muscle of swimming vertebrates forms a distinctive chevron morphology (Fig. 1B) (8, 44), which is believed to aid swimming (45–48). This structure is highly robust and symmetric between the left and right sides of the embryo (49). In zebrafish, myocyte progenitors for slow fibres remain mononucleated, whereas myocytes forming fast fibres undergo cell fusion to generate multinucleated cells (40, 41, 50, 51). Slow fibre myocyte progenitors differentiate into superficial slow fibres (SSFs), which line the lateral-most surface of the myotome, and muscle pioneers (MPs) that form the myoseptum at the dorsal-ventral midline (Fig. 1C) (41–43, 52, 53). The myocytes have complex morphodynamics, including elongation, fusion and coordinated rearrangements (40, 53, 54). Despite these significant morphological changes, the skeletal muscle forms with a robust internal structure, with closely packed and ordered myofibres by 72 hours post fertilisation (hpf). Therefore, myotome development offers an excellent system to explore how complex 3D cell morphodynamics with different cell types leads to an ordered structure at a tissue scale.

**Figure 1.**
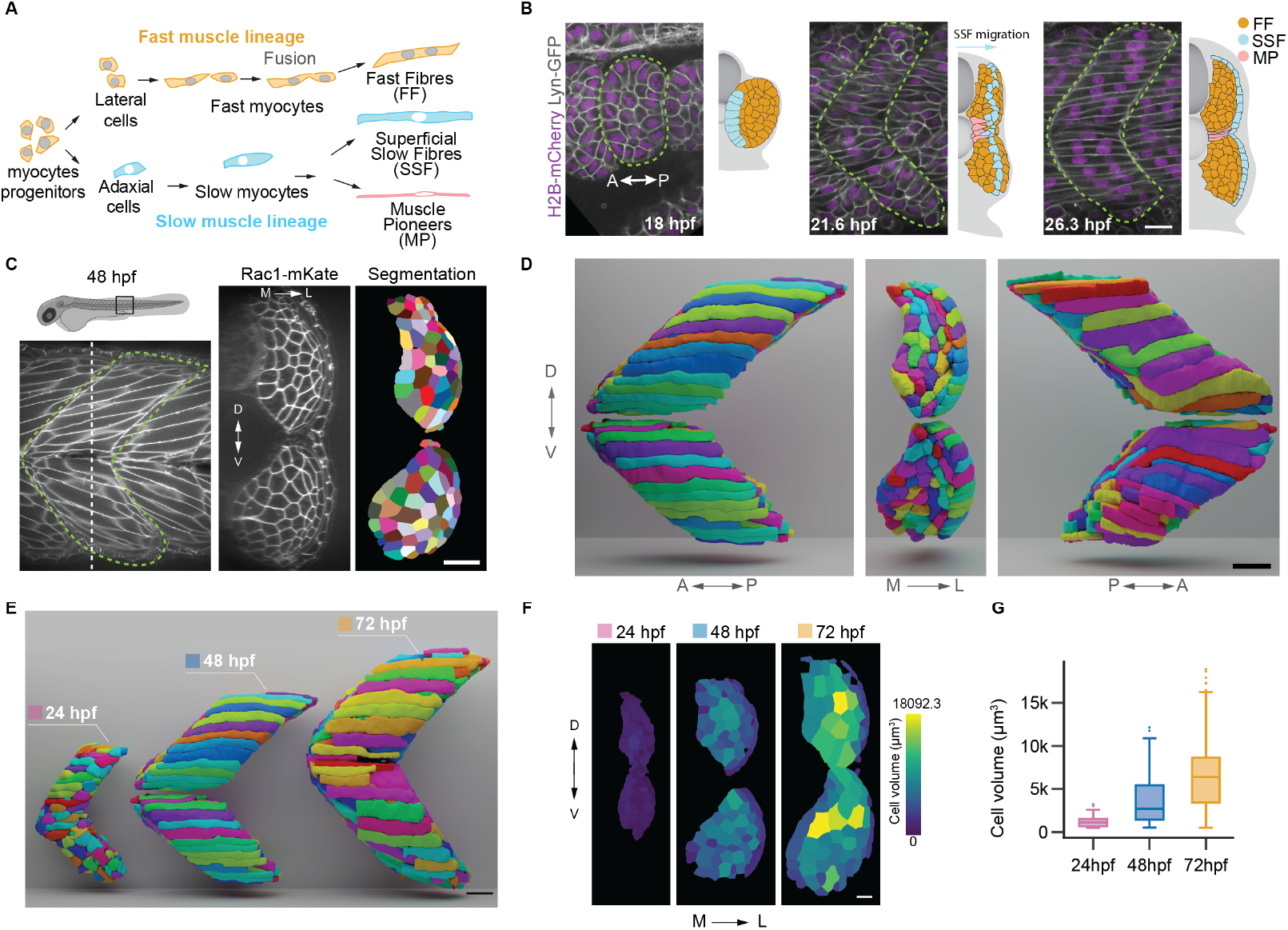
*in toto* muscle fibre analysis. **A)** Schematic of muscle cell populations. **B)** Snapshots of myotome formation from somite segmentation (18hpf) through to the formation of muscle fibres in an ordered array. **C)** Cross-sectional view showing imaging data (left) with cell segmentation on right at 48 hpf. **D)** 3D rendering of segmented myotome at 48 hpf under different views. **E)** Myotome rendering at different time points, plotted on same scale. **F)** Cell volume super-imposed on different time points. **G)** Cell volume quantification at different time points, *n* = 3. Scale bars 25*μm*.

We first developed an imaging and analysis pipeline to enable extraction of quantitative data about cell morphodynamics during the initial stages of skeletal muscle formation. We demonstrate that individual myocytes undergo coordinated twisting, which is mirror-symmetric between the dorsal-ventral and left-right domains. Further, we show that cell size, orientation and length varies significantly between different layers of the myotome, even between cells of the same type; each mature myotome segment has a distinct muscle configuration at a cellular scale but global patterns in structure (*e.g*., direction of cell twisting). Fast fibre myocyte progenitors are highly dynamic until they elongate to span the whole segment, when they undergo a rapid decrease in motility. By analysing cell packing and shape in mutants that alter cell fusion and specification along with biophysical perturbations, we propose that local cell shape plasticity is essential in ensuring robust myotome formation. This contrasts with typical epithelia, where cells are under high tension and ordered through long-range force propagation (16). These results are consistent with the developing muscle segments undergoing a structural transition, whereby the future fast myofibres freeze into a specific configuration once they span the myotome segment.

## Results

### Quantifying cell morphology during myogenesis

A long-standing challenge in organogenesis has been analysing cell shape in 3D within *in vivo* systems (55). With improvements in microscopy, we can now collect high spatiotemporal resolution imaging of the developing myotome (Fig. 1B, Movie S1, Methods). An advantage of using the developing skeletal muscle is that each muscle segment is defined in a sequential clock-like fashion (56, 57). Multi-position imaging enabled us to observe the whole vertebrate column, providing an effective time-course with approximately 30 mins between each segment (oldest segments most anterior), (Fig. S1A). Segmenting such data has been made substantially more efficient through machine-driven tools. We trained models within CellPose (Methods, (58)), fine-tuned for the different time points (*e.g*., 24, 48, 72 hpf), (Fig. 1C-E). From this segmentation, we visualised complete myotome segments in 3D using a custom environment within Blender (Fig. 1D-E, Movie S2, Methods). Our image analysis pipeline is freely available (59). To address the challenge of extracting 3D cell information, we utilised *CellMet*, a user-friendly tool that enables rapid and reliable extraction of the cell morphospace (60). We extracted morphological properties of the cells along with their connectivity and neighbour exchanges within the tissue.We make several initial observations, building on previous more qualitative work (39, 54, 61–63). First, the absolute size of each segment increased by around a factor of three from 24-72 hpf (Fig. 1E-G, Fig. S1B). This was driven by an increase in cell size, with myofibres being appreciably larger, rather than cell division (64). Second, the morphology of the slow and fast myofibres were distinct at 48 hpf. Superficial slow fibres were straighter, forming an ordered 2D layer (Fig. 1E). In contrast, the more medially-located fast myofibres took on a more diverse array of morphologies. There was clear change in cell size along the medial-lateral (ML)-axis (Fig. 1G right, Fig. S1B). Despite these variations at a cell scale, the tissue remained confluent throughout morphogenesis.

### Myocyte shape extends beyond cylindrical morphology

Within the myocyte population leading to fast myofibres, cell orientation relative to the embryo anterior-posterior (AP)-axis varied between medial and lateral domains (Fig. 2A). Near the centre of the myotome, cells aligned with the AP-axis (Fig. 2A(ii)). Yet, medial cells skewed outward, while lateral cells skewed inwards (Fig. 2A). We see that cell alignment is position-dependent within the myotome and mirrored across the dorsal and ventral domains (Fig. 2B). Observing the alignment over time, the distribution of cell orientations emerged during myogenesis as cells adapted to the physical environment; their initial elongation did not have such an angle distribution (Fig. S2A). Most fully elongated cells did not align either perpendicular to the boundary or with the AP-axis. This suggests that physical interactions between the actin-rich myotome boundaries (49) and the forming fibres do not determine the cell orientation.

**Figure 2.**
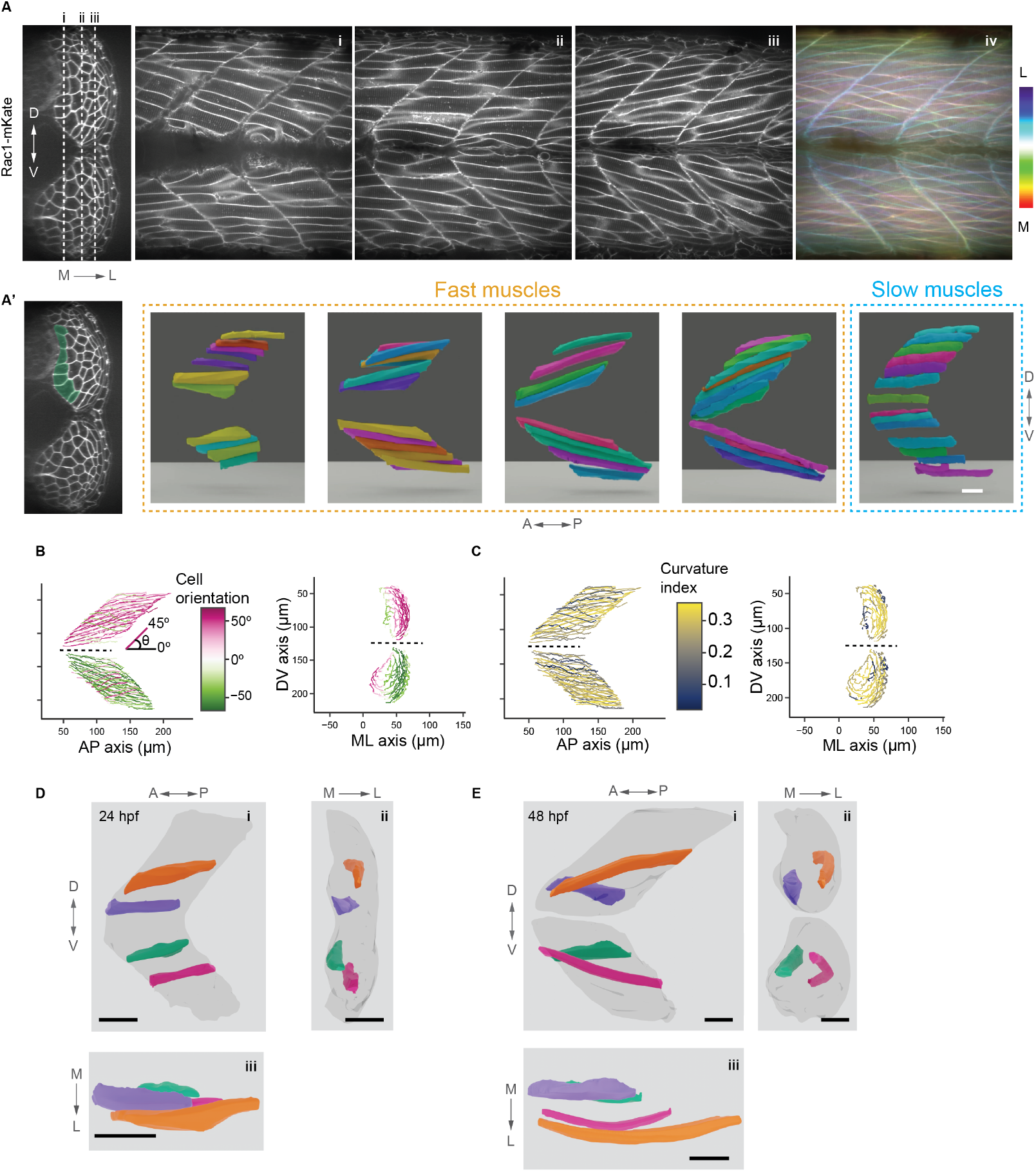
Cell shape during early myogenesis. **A)** Selected optical planes of embryo expressing Rac1-mKate: (i) medial-most layer of fast fibres; (ii) central layer of fast fibres; (iii) lateral-most layer of fast fibres at 48 hpf. (iv) Merger of all optical planes, colour coded by depth. **A’**) Cell rendering of slow and fast muscles grouped by layers through the myotome. **B)** Quantification of cell orientation relative to AP-axis at 48 hpf. **C)** Quantification of cell curvature at 48 hpf. (B) and (C) show one particular myotome segment; we see similar behaviour in three independent embryos. **D-E**) Selected cells in 24 hpf (D) and 48 hpf (E) embryos shown along different axes to highlight the onset of cell bending. Scale bars 25*μm*

Myofibres are typically portrayed as columnar-like, extending linearly between the anterior and posterior segment boundaries (65, 66). From our segmentation, we observed that cells, particularly near the medial surface, displayed a curved (bending) morphology (Fig. 2C, Fig. S2B). We used CellMet to quantify the cell curvature at 48hpf. We saw that the myofibres within the myotome bulk had highest curvature, with cells near the medial and lateral boundaries being straighter. In particular, the superficial slow fibres populating the most-medial section of the myotome segment were straightest.

To analyse this observation further, we compared cell shape from myotomes in similar AP position, in 24 and 48 hpf embryos (Fig. 2D-E). Early in myogenesis, the cells were straight (Fig. 2D), even after cell elongation was nearly complete. By 48hpf we see clear bending in cells (Fig. 2E), especially for cells towards the more lateral region. The cells displaying largest curvature were also tilted with respect to the AP axis (magenta and orange cells in Fig. 2E). Such curvature suggests these cells were not under high tension despite their elongated structure. The cell bending occurred after the initial phase of cell elongation, implying that the bending itself was not due to an intracellular process (*e.g*., polarisation in the cell cytoskeleton (67)).

We observed a population of myocytes that skewed relative to the AP-axis (Fig. 2E). These cells traversed multiple layers through the myotome. Such cell skew emerged as the cells elongated (Fig. 2D-E and Fig. S2); the skew may be due to space constraints as the cells elongated. Analysing the alignment at 48hpf, we saw that cells positioned towards the medial surface underwent the largest skew relative to the AP-axis (Fig. S2C). This corresponds to the region of the myotome with the greatest curvature.

Combined, these observations reveal that myofibre organisation within the myotome is highly complex, and not simply a case of columnar packing. Cell orientation and curvature both evolve with position and time, while maintaining a confluent environment.

### Muscle fibres undergo oriented twist during myogenesis

From our 3D segmentation of 48hpf embryos, we observed myocytes that twisted about the anterior-to-posterior axis (Fig. 3A, Movie S3). We used CellMet to quantify the rotation in the myocytes at 24, 48 and 72 hpf (Fig. 3B). Strikingly, the twisting was oriented, with mirror image behaviour between the dorsal and ventral domains at 48 hpf. The increase in twist correlated with the formation of the myoseptum at the DV-midline (Fig. 3B). The superficial slow fibres displayed a small twist, which was opposite to the bulk fast fibres. These twist domains were preserved to 72 hpf (Fig. 3C), with bimodal twist angle distribution.

**Figure 3.**
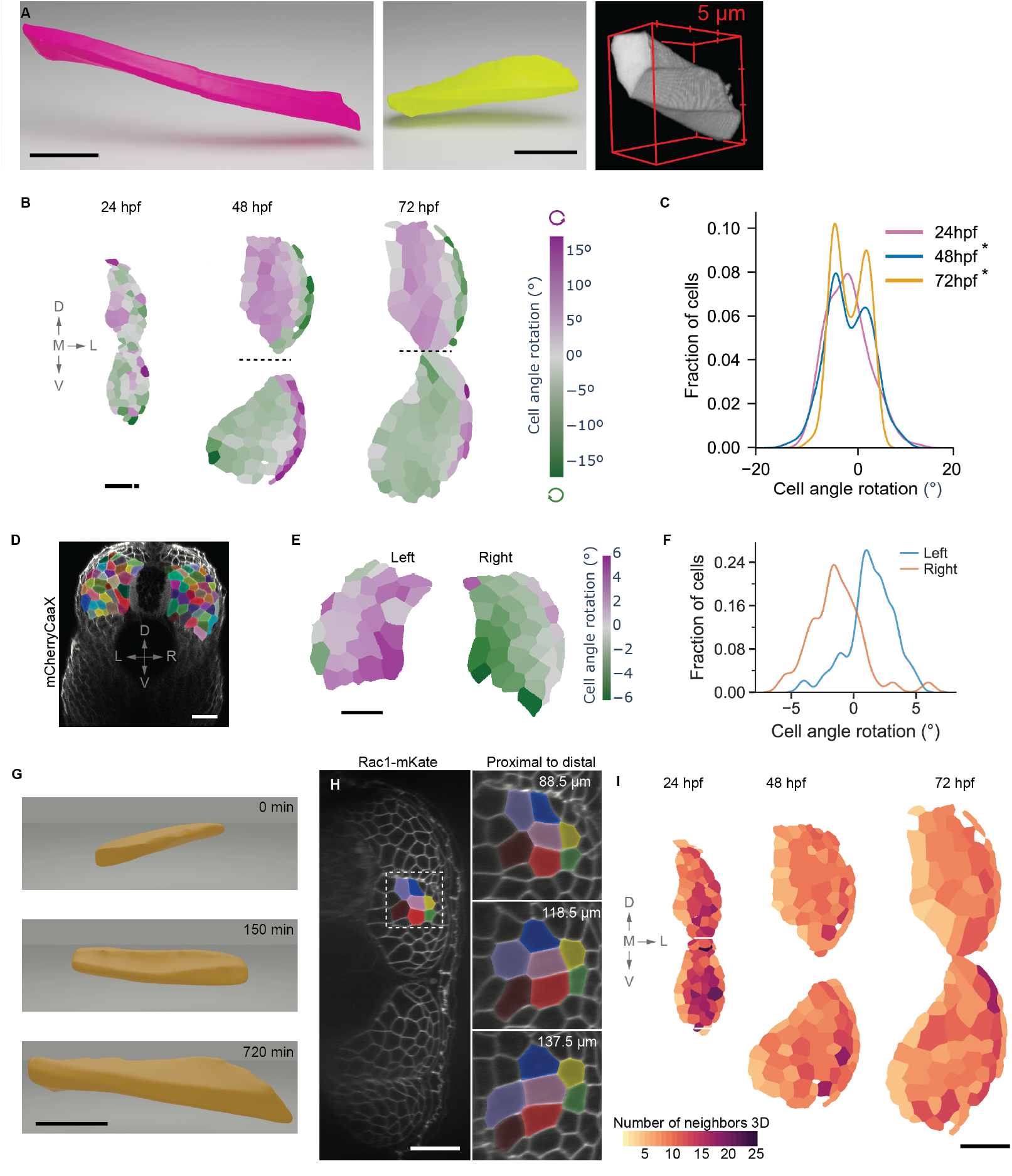
Muscle cell twisting. **A)** Example cell segmentation from lateral (left) and medial (centre) regions. Right shows a cell undergoing >45^°^ twist. B-C) Quantification of cell twist at different times. ^*^*p* < 0.005 Hartigans’ dip test for unimodality. Cross-section plots shows a representative myotome. *n* = 3 embryos per time. **D)** Cross-sectional view of a 72 hpf embryo, with CellPose segmentation overlaid. **E)** Extraction of cell twist in left and right sides of 48 hpf embryo. **F)** Quantification of cell twist between left and right sides of embryo (*n* = 3 embryos). **G)** Emergence of cell twist in a fast muscle myocyte during elongation. **H**) Analysis of neighbour exchange as cells twist in a somite stage 19 embryo. **I**) Quantification of neighbour exchange at different time points. Scale bars 25*μm*.

Given the above results, we hypothesised that the onset of cell twist is due to the curved boundary constraints. If this is the case, then the twist on the left-right sides of the embryo should be mirror-symmetric. We quantified images of 48 hpf embryos imaged such that both left and right sides are captured (Fig. 3D). Quantifying the cell twist, we saw mirror-symmetry in the angular rotation (Fig. 3E-F).

When does the twist initiate during myogenesis? We segmented 12 cells from our live movies every 30 minutes (Fig. 3G, Fig. S3A-B, Movie S4). We observed that the cells displayed a twisting morphology prior to adhering to both anterior and posterior domains of the myotome. This suggests that the twisting did not arise from torsional differences between the myotome boundaries along the AP-axis. Instead, the twisting was an emergent property of the cells as they elongated.

Though we see that there is global order in the twist, the amount that each individual cell twisted was variable (Fig. 3H). If neighbouring cells twist at similar amounts (*e.g*., like threads wound to make a rope), then the neighbour number should remain similar along the cell length. However, we did not see such behaviour when looking at groups of neighbouring cells (Fig. 3H), with significant cell-to-cell variability apparent, even at similar locations. We further traced the position of the cell centre along its anterior-to-posterior axis relative to neighbours, and we saw no clear local ordering in the twist (Fig. S3C).

We extended this analysis to myotome segments at 24, 48 and 72 hpf (Fig. 3I). We quantified average neighbour number along the whole cell length. If the system was perfectly columnar, the neighbour number would be six (excluding myotome boundaries). At 24hpf, when cells were still roughly spherical the mean number of neighbours was around 10-12, (Fig. S3D) (perfect sphere packing would be 12 neighbours). As the cells elongated, the packing fraction reduced, consistent with sphere-to-columnar morphological changes. The mean packing fraction was 8.5 ± 2.0 at 72 hpf (Fig. S3D). Though smaller than sphere packing, this was still substantially larger than expected for columnar packing. To conclude this section, we see that there are clear patterns in the twisting of fast fibres across the myotome. Yet, there is substantial variability in the measured twist at a single cell scale, even for cells at similar location.

### Cell shape transitions during myogenesis

Myotome formation is morphologically complex, with cell skew, bending, fusion, and twisting. To analyse the morphology between different stages, we next turned to a spherical harmonic representation of the cells (Methods, Fig. 4A) (68). We performed a statistical clustering of the spherical harmonics to reduce the complexity (Methods). Our approach enables users to interrogate specific cells within the statistical clustering and map back onto the morphological position (Movie S5) (59). We saw that at 48 hpf, cell groups - such as superficial slow fibres and medial lying fast fibres - formed distinct clusters within this reduced representation (Fig. 4B).

**Figure 4.**
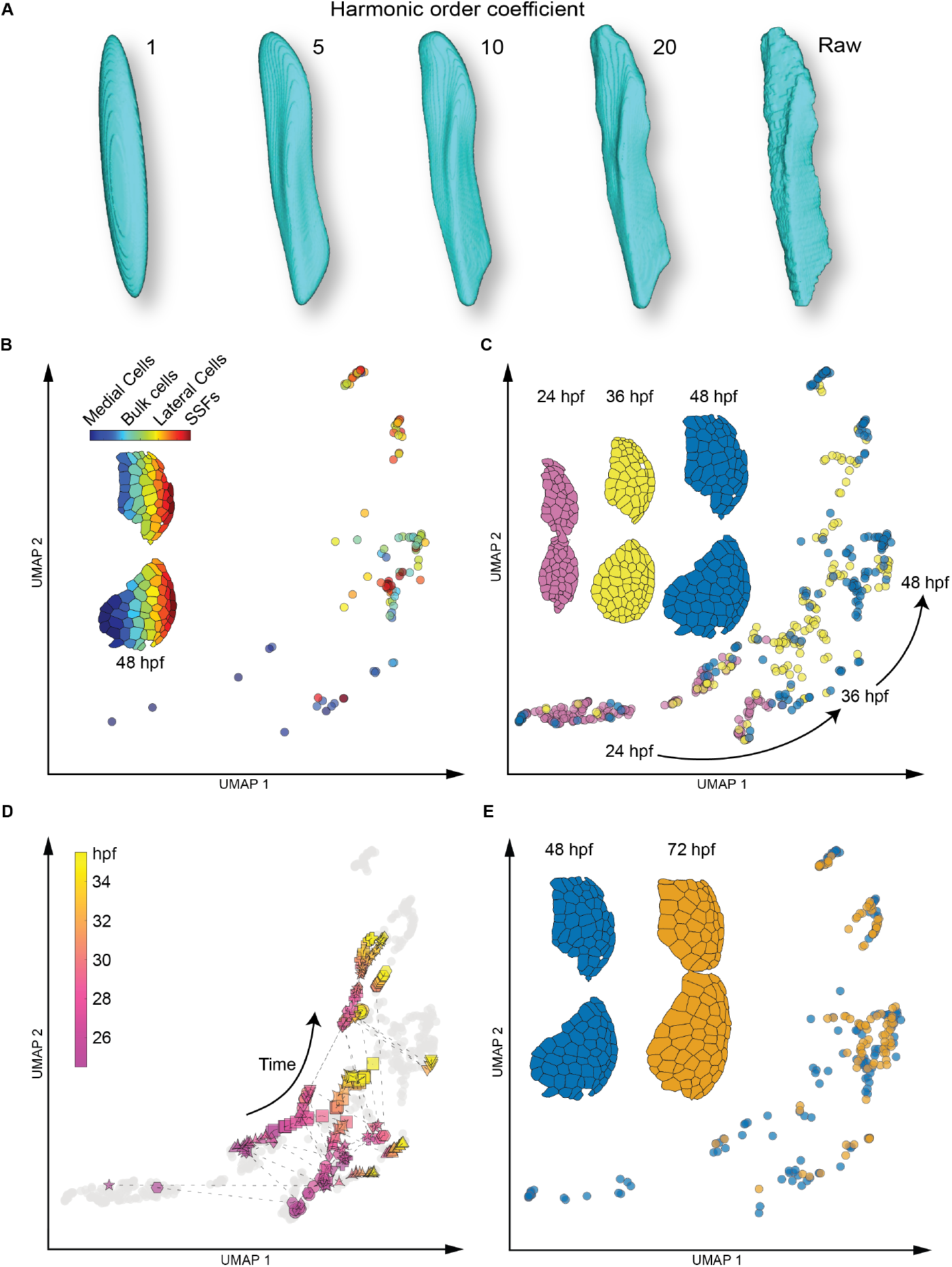
Statistical analysis of cell morphometrics. **A)** Outline of approach to represent cells in terms of spherical harmonics. **B)** UMAP of spherical harmonic parameters at 48 hpf with corresponding cross section of myotome segment with cells labelled by medial-lateral position (Methods). **C)** UMAP of 24, 36 and 48 hpf myotome segments, each from a different embryo, with corresponding cross sections of myotome segments. **D)** UMAP of the trajectories of 10 cells from 24 to 36 hpf in 30 min intervals. Dashed lines represent trajectories and different symbols correspond to specific cells. Colour coding represents time from pink (initial) to yellow (final) time points. **E)** UMAP of 48 and 72 hpf segments with corresponding cross section of myotome segment.

Between 24 to 48 hpf there was a clear shift in the morphology space (Fig. 4C); cells at 24 hpf and 48 hpf were largely distinct. This is consistent with the approximately spherical cells at 24 hpf becoming elongated by 48 hpf. The segment at 36 hpf lies near the transition point between the spherical and cylindrical structural arrangements, with subsets of cells lying either close to 24 or 48 hpf morphologies.

To better understand these dynamics, we followed the behaviour of 10 cells segmented throughout their initial elongation phase (Fig. 4D). These cells followed a similar general trend through the morphological space. But, at a single cell level there were distinct paths, with different final positions within the morphological space (Movie S6). This is consistent with the above observations that at a tissue-scale myotome segments are highly similar but there is large cell-to-cell scale differences both within and between myotome segments. This result tallies with previously observed stochasticity in the timing and positioning of muscle fusion (40).

We next asked whether the morphological space further altered from 48 to 72 hpf. Surprisingly, there was little change in the geometric structure of the myotome segments (Fig. 4E), despite the 72 hpf segments being physically larger. This suggests that the structure of the segments was fixed at 48 hpf, with only simple growth dictating the change to the morphology at 72 hpf. These results are consistent with the cells freezing into place by 48 hpf; the specific cell configuration varies between segments, but once fixed in place within a segment the cells remain largely unchanged morphologically.

### Altering cell shape and fate impacts the material behaviour

We explored cell behaviour under two perturbations: (i) disruption of cell fusion; and (ii) inhibition of slow muscle cell fate. Inhibiting cell fusion using a *mymk* mutant (69) led to thinner cells (Fig. 5A(ii)). Essentially, the number of cells nearly doubles though the nuclei number remains similar. *mymk* mutant embryos still displayed cell twisting at 48 hpf (Fig. 5B, Fig. S4A-B). However, the twist was noisier, with less clear differentiation in twist between slow and fast fibres (Fig. 5B). *mymk* mutant embryos lost the clear double-peak in twist angle observed in wildtype embryos (Fig. 5C).

We next assayed the cell shape and alignment in *mymk* mutant embryos. The cells underwent neighbour exchanges along their extent, in part due to them displaying a crinkled morphology (Fig. 5A(ii), Fig. S4B). The cell alignment with the AP-axis was largely unaffected (Fig. 5D). There were no apparent extracellular spaces in the myotome and we note that *mymk* mutant zebrafish are viable as adults, though with decreased swimming capability (69).

How is the myotome structural order affected by perturbation to the cell state? We examined the cellular morphospace in *uboot* mutants (62) (Fig. 5A(iii)), which do not have slow muscle fibre populations - all cells in the myotome become multinucleated fast fibres. Myofibres in *uboot* mutant embryos mostly remained untwisted (Fig. 5C, Fig. S4C). Further, the bimodal cell alignment distribution was lost; the majority of cells were aligned with the AP-axis (Fig. 5D). Despite these morphological changes, the segments still maintained structural integrity at 48 hpf, with no significant extracellular spaces apparent.

**Figure 5.**
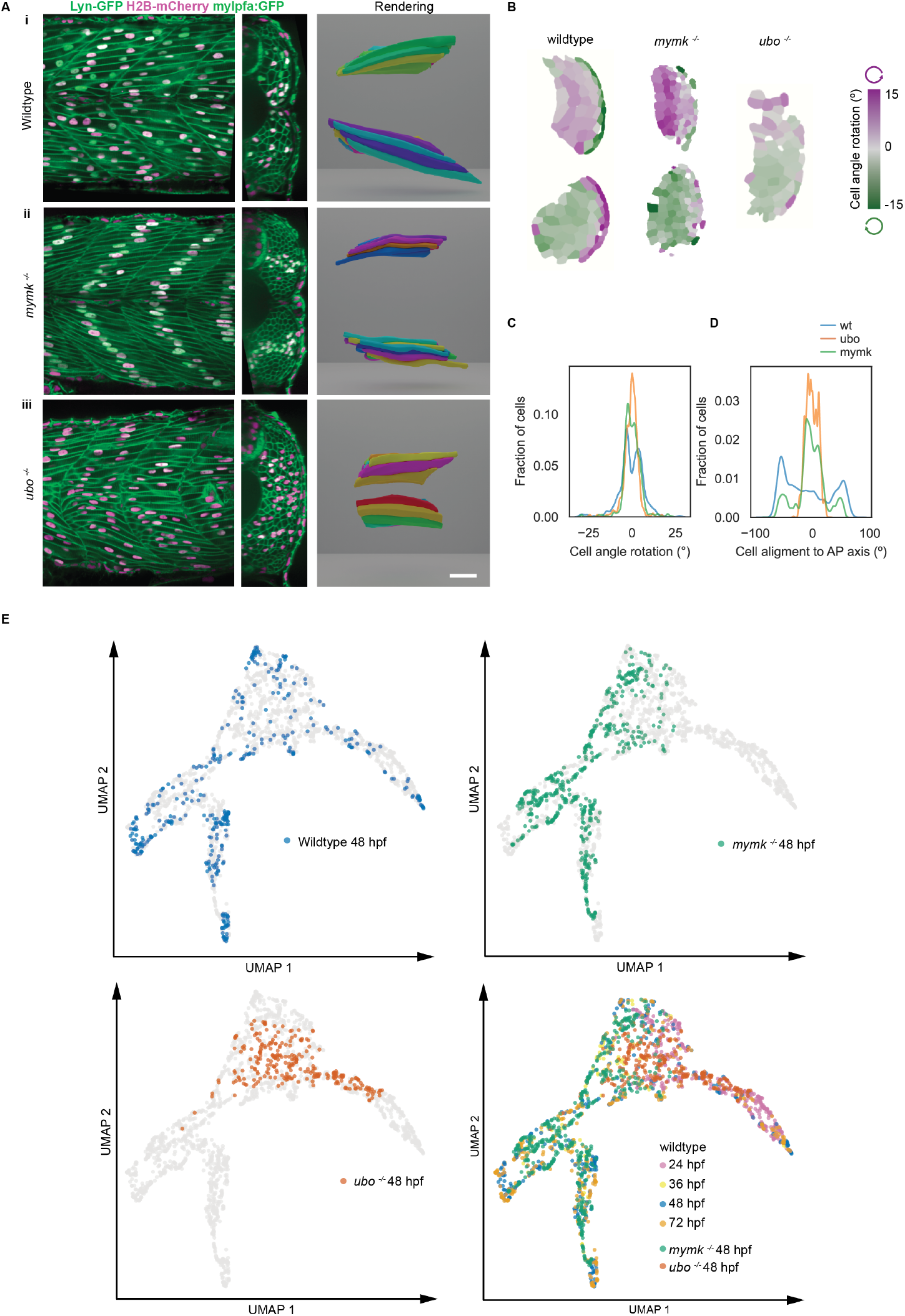
Perturbing cell morphogenesis and fate. **A)** Left column: Wildtype (i), *mymk-/-* and *uboot-/-* embryos at 48 hpf. Central column: cross-sectional view through the myotome with the myoseptum (or absence thereof) highlighted by dashed line. Right column: example cell rendering from similar locations within the myotome. **B)** Cross-sectional view of cell twisting. **C)** Quantification of cell twisting at 48 hpf in the different conditions (*n* = 3). **D)** Distribution of cells aligned to the AP-axis at 48 hpf (*n* = 3).E) UMAP of spherical harmonic representation comparing cell shape in the different genotypes at 48 hpf. (WT 24hpf: n=2; WT 36hpf: n=1; WT 48hpf: n=3; WT 72hpf: n=3; mymk^-/-^ 48hpf: n=3; ubo^-/-^ 48hpf: n=3)

To further test how these perturbations impacted cell morphology, we mapped the spherical harmonic representation of 48 hpf embryos under different conditions (Fig. 5E). There were two notable behaviours. First, the cell shapes of the *mymk* and *uboot* mutants lay within the shape space defined by wildtype embryos. This is consistent with these perturbations still leading to confluent tissues with elongated fibres. Secondly, the *mymk* and *uboot* mutants populated specific subspaces of the morphological space. In particular *uboot* embryos only populated a small part of the available morphological space.

### Cell dynamics shift during fibre elongation

Along with the changes in cell shape, cells are also highly dynamic during the period of initial muscle formation (40). How do cell dynamics change during myogenesis?

We tracked 474 nuclei ((40) for details) during myotome formation (Movie S7). In wildtype embryos, we noticed a sharp decrease in motility around five hours post somite segmentation (Fig. 6A). This rapid decrease in nuclei speed correlated with the elongation of cells spanning the segment along the AP axis (Fig. 6A, Fig. S5A).

**Figure 6.**
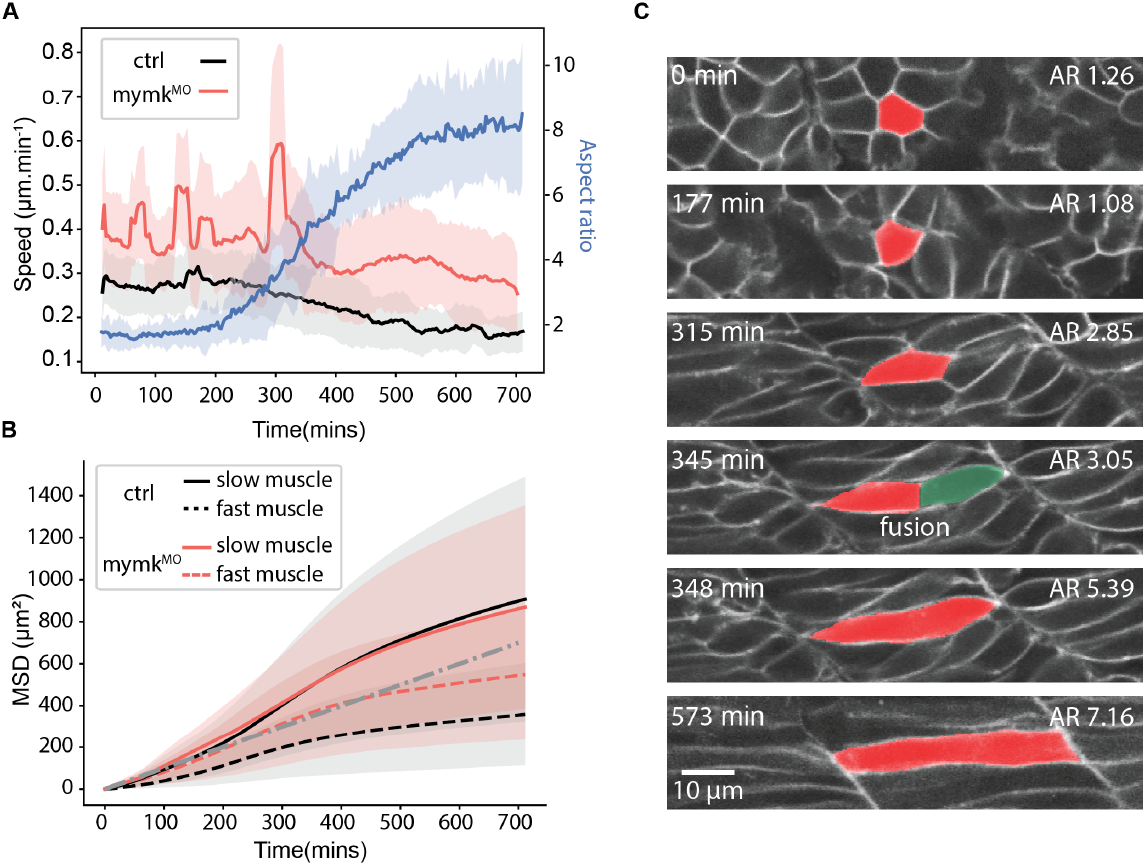
Cell dynamics during myotome formation. **A)** Nucleus speed with respect to time after somite segmentation. ctrl: 474 tracks from *n* = 2 myotomes. mymk 193 tracks from *n* = 2 myotomes. Right axis corresponds to cell aspect ratio of wild type cells (*n* = 17 cells from 2 myotome segments). Shading represents ±1 s.d. **B)** Mean-squared-displacement for wildtype (ctrl) and *mymk*^*MO*^ embryos, separated into fast (dashed) or slow (solid) muscle fibre progenitors. Dashed grey line represents *α* = 1. **C)** Example showing how cell fusion leads to rapid cell elongation in a wildtype embryo expressing Lyn-GFP. Fusion occurs at 345-348 min post somite segmentation. The aspect ratio (AR) is shown in the upper right corner.

To test this more rigorously, we measured the nuclei mean-square-displacement (MSD). The scaling of MSD with time (determined by an exponent *α*) reveals information about the *∼*system dynamics, *MSD t*^*α*^. *α* = 1 corresponds to a simple diffusive process (70). Prior to fusion, the MSD exponent *α* for myocytes that became fast myofibres was between 0.5 and 1.0; consistent with a restricted diffusion process (Fig. 6B, Fig. S5B). Post fusion, the steepness of the MSD decreased, leading to *α <* 0.5. The displacement post fusion was largely due to nuclei repositioning within the fused cells, and not movement of the cells themselves (40). The future slow muscle myocytes had a large MSD gradient (*α >* 1.0) during early myotome development (Fig. 6B). This was consistent with the directed movement of the future superficial slow fibres to the medial layer. We saw a distinct decrease in the MSD gradient once the cells spanned the myotome segment. As with the future fast fibres, the remaining contribution to the MSD appeared to come from nuclei repositioning within the cell, and not cell movement.

To live image cell behaviour under perturbation of fusion, we took advantage of the well-characterised *mymk* morpholino (Fig. 6A, Movie S7). In this perturbation, cells completed elongation along the AP-axis more slowly; in effect, fusion of two partially elongated cells in wildtype embryos often led to one elongated cell and this process was absent in *mymk*^*MO*^ embryos (Fig. 6C). This suggests that while fusion is not necessary for eventual elongation, it is required for rapid freezing of the muscle configuration into a rigid state. In accordance with this conclusion, the dynamics of the future slow fibres remained largely unchanged in *mymk*^*MO*^ embryos, with no significant difference in the MSD behviour to wildtype embryos (Fig. 6B).

The cell morphodynamics during the initial phase of myotome formation (Figs. 4-6) are consistent with the myotome segments undergoing a structural transition during morphogenesis (20). This transition represents moving from a fluid-like state, where cells can change shape and position relatively freely (*α >* 0.5), to a frozen solid-like state where cell movement and rearrangement is hindered (*α <* 0.25).

### Myocytes are not under tension and mechanically malleable

Our above conclusion is dependent on the cells being able to mechanical adapt to the changing conditions during early myogenesis in order to freeze into a specific configuration while leaving no appreciable extracellular space. Next, we focused on understanding the mechanical state of the myocytes and how they mechanically respond to perturbations.

We first examined the distribution of actin during myotome maturation (Fig. 7A). Coordinated multicellular actin structures can drive organogenesis, such as in the heart (71). However, we did not observe evidence for such structures in the developing myotome. Instead, as previously reported (50), we observed actin localised to subcellular puncta during fusion. This suggests that long distance actin-driven processes are not shaping the tissue.

**Figure 7.**
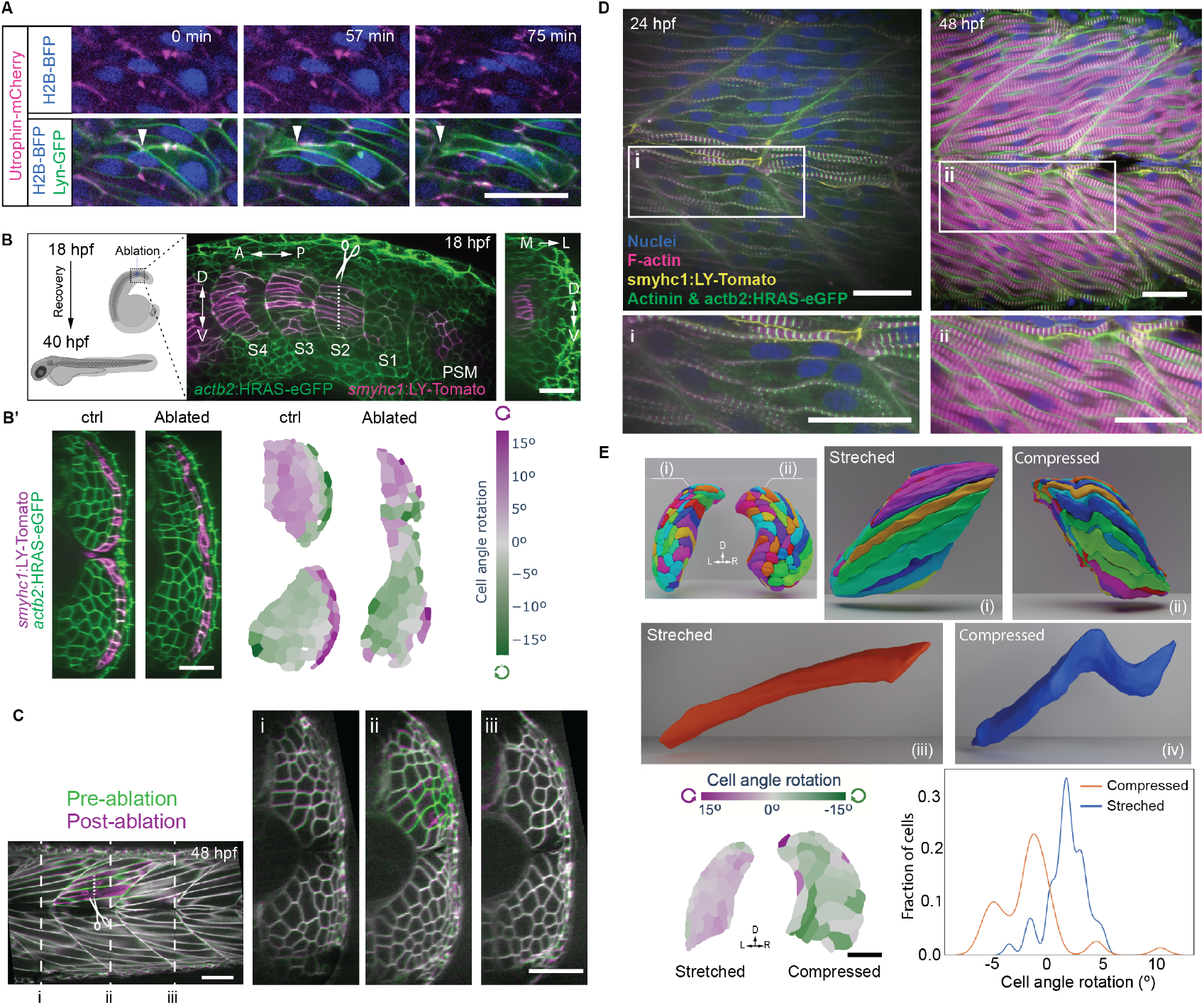
Developing muscle fibres are mechanically malleable. **A)** F-actin (Uthrophin-mCherry, magenta) localisation during fast muscle fibre elongation. Cell membrane (Lyn-GFP, green) and nucleus marked (H2B-tagBFP, blue). Arrow heads marks tip of elongating cell. **B)** Ablation of early elongated future muscle pioneers at 18 hpf affects horizontal myoseptum formation. Cells express actb2:HRAS-GFP (cell membranes, green) and smyhc:LY-Tomato (slow muscle reporter, magenta). Cross-section at the ablation site is shown. **B’)** Left. Myotome structure without or with laser ablation of horizontal myoseptum. Right column shows quantification of cell rotation at 48 hpf. **C)** Ablation of myocytes during elongation. (Left) Overlay of tissue before and 5 minutes after ablation. (Right) Cross-sectional view of myotome segment before and after segmentation at the mytome ablated and neighbouring myotomes. **D)** Actinin localisation (green) at 24 hpf (left) and 48 hpf (right) counterstained with DAPI (blue), phalloidin (magenta) and combined with expression of smyhc:LY-Tomato (yellow, slow muscles reporter) and cell membranes (green, actb2:HRAS-eGFP). **E)** Rendering of myotome segments under biophysical perturbation (top row), with example cells shown below. Data from (73). (Bottom): Analysis of cellular twist under different confinement conditions in a representative embryo. Scale bars 25*μ*m.

To more directly assay the tissue mechanical properties we used laser ablation at different stages during myotome formation. We separated our analysis into two targets: (1) the forming muscle pioneers which appear to act as an important scaffold for the developing myotome; and (2) the extending myocytes in the bulk. Ablating muscle pioneers before myocyte elongation resulted in the loss of the myoseptum (Fig. 7B-B’ left, Movie S8). Yet, the tissue restored confluency, suggesting the myocytes can adapt to local mechanical perturbation. In this case, we saw a reduction in the cell twist (Fig. 7B right). A major structural difference between wildtype and *uboot* embryos is the absence of the myoseptum in the latter case (Fig. 5A(iii)). Combining our results from the mutant and ablation analysis, we posit that formation of the myoseptum generates constraints that, at least partially, induces cell twisting and skew in wildtype embryos, as the cells maintain confluency while also elongating into myofibres.

Ablating muscle pioneers after the myotubes had formed resulted in little structural change, even though they were clearly force bearing (Fig. S6A, Movie S9). This supports the observation that once the fast myofibres form, they are frozen into a particular configuration. Finally, we ablated myocyte progenitors for fast myofibres during the elongation phase (Fig. 7C, Movie S10). We noticed little change in neighbouring cell morphology or position, consistent with these cells not being under significant mechanical load or generating large internal forces (Fig. S6B). Further, cells that had fully elongated appeared to not change shape or position after the ablation. This suggests the myocytes themselves are not the primary factor actively driving the forces reshaping the whole segment.

Given the cell malleability, what is the process driving the rigidity of the cells once they span the myotome? Recent *in vitro* work has revealed that myocytes are highly dynamic until regular sarcomeric structures form. Once sarcomeres are apparent, cell motility rapidly decreases (72), driven by long-ranged tension. These observations are consistent with our ablations; if we ablate after myofibre structures have formed, we observe recoil but not if cells are still elongating. To further test this, we performed antibody staining for sarcomeric structures at 24 and 48 hpf (Fig. 6D). At 24 hpf, we see the emergence of sarcomeric structures in the muscle pioneer population (Fig. 6D(i)). The fast muscle population at this stage has lower expression of actinin and no apparent intracellular actin structures. In contrast, at 48 hpf, when the cells are (largely) fixed into place, we see clear sarcomeric structures throughout all muscle fibres, including repeated actin domains (Fig. 6D(ii), Fig. S1A inset). This suggests that formation of tensile sarcomeric structures underlies the maintenance (but not formation) of the ordered muscle state.

### Boundary constraints guide muscle shaping and consequently myotome morphology

To further test our hypothesis that boundary constraints are important in driving cell morphodynamics in the developing myotome, we considered two further perturbations: biomechanical and genetic. Using actively trapped embryos, the developing myotome can be subjected to different external constraints. Under trapping, the myotome on one side is compressed, while the opposite side is stretched or elongated (73). We ran our analysis pipeline on these deformed myotome segments and quantified the resulting cell shapes (Fig. 7E). We saw that cells buckled, yet maintained confluent integrity of the myotome (Fig. 7E(iii)).

We quantified the cell twist in embryos which were partially constrained. The stretched side displayed similar behaviour to wildtype (Fig. 7E), with a biased twist. The compressed side showed much larger variability in the cell twist. This is consistent with the cell shapes adapting to the changing boundary constraints. Despite these morphological variations, we did not observe any significant extracellular space within the myotome at any stage from 24-72 hpf.

An alternative way to disrupt boundaries is genetic. We used *fused somite* embryos, which do not form clear boundaries between somite segments (74). Myofibres still formed but in a more disordered fashion (Fig. S6C). We saw myocyte elongation in some cases perpendicular to the AP-axis. Further, in 4 out of 7 movies, we observed the loss of tissue confluency (Fig. S6D). In segments that maintained confluency, we saw large variability in both cell morphology and slow muscle position (Fig. S6E). The loss of ordered cellular packing correlates with decreased robustness in tissue-scale morphogenesis.

## Discussion

Organ development requires multiple steps, from cell fate determination and cell shaping through to whole tissue reconfiguration. Yet, this process happens with remarkable robustness. Here, we demonstrate that skeletal muscle forms robustly by undergoing a structural transition, whereby motile fast fibre myocytes are locked into place once they span each myotome segment. To facilitate this spanning, the cells are mechanically malleable, enabling them to twist and skew through the dense cellular environment as they elongate. Cell-cell fusion enhances the rate of elongation, as the fusion of two partially elongated cells typically leads to one (near to) fully elongated cell. These morphological changes, while noisy at a single cell scale (40), display long-ranged order (*e.g*., ordered twist) due to the specific boundary constraints imposed during morphogenesis. The cell length extent along the AP-axis may act as the effective order parameter, with the myotome segment rigidifying once the majority of cells span the segment.

This work reveals how multiple cell types can interact in 3D to build a fully confluent tissue. Future slow fibres are the first to elongate and span the myotome compartment. The muscle pioneers form the myoseptum, defining a central, curved domain under tension. Meanwhile, the superficial slow fibres migrate to the myotome lateral surface where they form a rigid boundary. Disruption to these boundary-defining cells (either through mutants or ablation) perturbs cell packing. Boundary constraints have been shown to play a role in cell confinement (75), vortex pattern emergence (76, 77), response to wounding (78) and tissue shaping (8). Previous work has highlighted how cell shape impacts jamming transitions in effectively 2D biological systems (79). Here, we have revealed that complex 3D tissues can undergo an effective structural transition when the cells have an ordered elongation and defined boundaries.

There is a correlation between the emergence of sarcomeres and the rigidity of the myotome (Fig. 6D). If the sarcomeres are functionally important in stabilising the myotome structure, then we would predict that disrupting sarcomere formation, *e.g*. through disruption of actinin or transplanting between wild type and mutant embryos (80), would impact the architecture of the forming myotome. In particular, there may be cell rearrangements or cell reshaping after the cell elongation process is complete.

We have not considered the role of cell adhesion. The adhesive properties of the cellular microenvironment - both to other cells and the surrounding extracellular matrix (ECM) - can play an important role in determining the tissue material properties (17, 81–83). Disruption of the ECM leads to rapid loss of myotome structure (8). During myogenesis there are known spatiotemporal changes in M- and N-Cadherin localisation (84), which may impact the cell motility and hence the structural transition. It will be interesting to re-explore these results with the more quantitative tools now available, to dissect the role of the ECM and adhesion molecules in myotome morphogenesis. Relatedly, it may be possible utilising imbedded magnetic beads to estimate the magnitude and extent of forces on cells from the boundaries (85) and how these evolve over time during morphogenesis.

Our work has focused on structural formation of the myotome during zebrafish embryogenesis. Is this structure maintained through to adult stages? Qualitative analysis of adult zebrafish embryos suggests that the fibres maintain a twisted morphology (45, 48). Modelling of such muscle structure suggests that this twisting may be beneficial for swimming efficiency in zebrafish (47, 48). It will be interesting to perform imaging across multiple stages of zebrafish growth to see whether the general myotome structure is preserved from embryo to adult. A second important question is whether such structures are evolutionarily conserved? Similar muscle structures have been observed in other fish (46), though not at cellular scale early in development. We investigated *A. calliptera*, a teleost 220 million years diverged from zebrafish. At 28 somite stage, cells in the myotome display twisted cell morphologies, consistent with our above results (Fig. S7A). We also examined cell structural order in segments of the closely related species *R. sp. chillingali*. We imaged embryos at the end of segmentation (Fig. S7B). We saw clear variation in alignment of fibres along the ML-axis, consistent with zebrafish. These observations suggest our results are broadly relevant to teleost development, though further quantitative work, especially in live embryos, is required to test this hypothesis.

Biological systems display a remarkable ability to self-organise to form complex tissues. Here, we have demonstrated the interplay of elongating future myofibres with tissue boundaries drives a structural transition that freezes cells into place, ensuring robust formation of the vertebrate muscle segments. During these stages, cells are mechanically malleable enabling them to adjust for local variations in cell size, cell neighbours, and tissue boundaries. This results in the unique configuration of each myotome segment at a cell scale. Our work opens new avenues into understanding how organs reliably emerge during development.

## Author Contributions

M.A.M-S and T.E.S conceived the project. S.T. and M.A.M-S developed the software for image analysis and visualisation. Y.H. developed the software for spherical harmonic representation. M.A.M-S performed most of the experiments. T.E.H. and R.G.P. performed the experiments physically constraining the developing embryo. S.E.T. and B.V. performed the collection and imaging of *R. sp. chillingal* and *A. calliptera*. S.T., M.A.M-S and Y.H. performed the data analysis. S.T. and M.A.M-S performed data visualisation. M.A.M-S and T.E.S wrote the first draft of the manuscript, with all authors contributing to the final submitted version.

## Material and methods

### Zebrafish husbandry

All zebrafish strains were maintained according to standard procedures for fish husbandry at the zebrafish facilities of the Institute of Molecular and Cell Biology, A*Star, Singapore and University of Warwick. The following wild-type, mutant and transgenic strains were used: AB, inbred wild-type control, *ubo*^*tp*39^ (86), *mymk* (69), *Tg(smyhc1:lyn-tdTomato)* (87), and *Tg(prdm1a:GFP)* (88). All experiments with the zebrafish in Singapore were approved by the Singapore National Advisory Committee on Laboratory Animal Research. All experiments in Warwick were performed in compliance with the University of Warwick animal welfare and ethical review board (AWERB) and the UK home office animal welfare regulations, covered by the UK Home office licenses PEL 30/2308 and X59628BFC to the University of Warwick.

### Live imaging and laser-ablation

Wild-type or mutant zebrafish embryos were injected with *h2A-tagBFP* and *lyn-gfp* mRNA for labelling nuclei and cell membranes respectively. Injected embryos were incubated at 25°C or 28°C until they were visualised. Embryos were mounted in low melting agarose. For immobilisation of embryos, embryo medium containing 0.003% tricaine or α*-bungarotoxin* mRNA injection at one-cell stage (89) were used. Mounted embryos were visualised using an Olympus spinning disk confocal system equipped with an inverted motorized microscope IX83, 60X silicon immersion objective (UPLASAPO60XS2, NA: 1.30, W.D: 0.30mm), a confocal scanning unit W1 (Yokogawa), two ORCA-Fusion BT CMOS cameras (Hamamatsu) and OBIS LAX lasers 405 nm, 488 nm, 561 nm and 640 nm (Coherent). Single volumetric stacks of whole mounted embryos were acquired using 200 ms exposure, 2034 x 2304 pixels image size, a 0.5 µm z-step size and 10% laser power. For time-lapse imaging, confocal images stacks were acquired every 30 min using same imaging conditions. Embryos were kept at 28.5°C using an on-stage incubator (Okolab). For laser-ablation experiments, a 355 nm pulsed laser was used (UGA-42 Caliburn, RAAP OptoElectronic) coupled to the Olympus spinning disk confocal system equipped with a 60x objective described above.

### Image pre-processing for cell segmentation

Due to the elongated phenotype of muscle cells, confocal stack images were resliced to generate stacks along the YZ planes.

### Deep learning single cell segmentation

Single cell segmentation was performed using the Cellpose 2.0 algorithm (58). Cellpose was run using the cyto2 model on a sample of 10 resliced images. The resulted masks were manually corrected using the Cellpose GUI. Cell contours outside the myotome region were removed for improving detection of the muscle cells. Manually edited mask images contained around 100 annotations per image and these were used for generating an intermediate fine-tuned cyto2 model. The fine-tuned cyto2 model was used to predict the cell contours on a sample of 50 images from 4 embryos. The labels obtained then were manually edited and used for training a model from scratch (300 epochs, 0.1 learning curve). The final model was used for prediction of volumetric stacks creating 2D labels on each XY slice and then stitching these across slices using the following parameters (diameter=140, flow threshold=0.4, cell-prob_threshold=0, stitch_threshold=0.2). Labels were saved as a multi-page tif files for post-processing.

### Cell segmentation post-processing

For ensuring only labels corresponding to muscle cells in the myotome were included, labels were measured using the Analyse regions 3D command from MorpholibJ (90) in Fiji (91). The obtained measurements (centroid, volume and surface) were used for filtering and preserve the labels corresponding to muscle cells in the myotome, labels with a volume size below 100k pixels or 5 × 10^3^*µm*^3^ and located outside the myotome were not included. After filtering, the centroids of the muscle cell labels were clustered and grouped to the corresponding myotome using the DBSCAN algorithm in R (92). For DBSCAN clustering, the optimal value for *E* was calculated for every full stack. Finally, muscle cells labels of single mytomes were saved as multi-page tif files.

### Three-dimensional visualisation of muscle cells at single cell resolution

For quick inspection of 3D volumes, the images files containing the labels were visualised using the 3D viewer in Fiji (93). For generating high detailed 3D renderings of cells, volumetric files containing the muscle cells labels were used for generating the corresponding polygon meshes using the 3D viewer in Fiji or the Meshio library in Python (94). Polygon meshes corresponding to individual cells were exported as single OBJ files and then imported to Blender v3.4.1 using the batch import OBJ plugin. For presentation purposes, polygon meshes were simplified by decimation (decimate modifier 0.1) and smoothed (smooth modifier iterations 99) in Blender. For generating still images and animations, meshes included in the scene were rendered using the render engine Cycles (noise threshold 0.1, min samples 0, max sample 1024). Animation renderings were exported as MP4 movie files.

### Spherical harmonic representation and statistical mapping

3D segmented images were separated into binary tiff files containing segmentations of each individual cell volume. SPHARM-RPDM (95, 96) was used to compute the first five degrees of spherical harmonic coefficients of each cell, for a total of 36 coefficients further split into x, y, z and real, imaginary components, giving 216 parameters. UMAP (97) was conducted using parameters *n*_*neighbours*_ = 15, *min*_*dist*_ = 0.1. Holoviews (98) was used to generate the interactive plots linking UMAP embedding to shape and positional information of the cell within a 2D cross section of myotome.

### Whole mount immunofluorescence in zebrafish embryos

Embryos were manually dechorionated and fixed in 4% formaldehyde for 2 hours at room temperature. After fixation, embryos were washed three times with PBST (0.1 Tween-20 in PBS 1X). Embryos were permeabilised and blocked for 4 hours at room temperature in blocking solution (1% Triton-X100, 1% DMSO, 1% BSA in 1x PBS). Embryos were incubated overnight at 4ºC in mouse anti-actinin antibody (A7732, Sigma-Aldrich) diluted 1:500 in blocking solution. Then, the embryos were incubated half day at 4ºC in goat anti-mouse Alexa 488 (A11001, Invitrogen) diluted 1:800 in blocking solution. Embryos were then stained overnight at 4ºC with phalloidin Alexa 647 diluted 1:100 in blocking solution. Finally, embryos were stained 1 hour at room temperature with DAPI (D9542, Sigma-Aldrich) diluted 1:10,000 in PBST. Embryos were mounted in 1% low melting point agarose diluted in PBS and imaged using a Olympus spinning disk confocal system equipped with a 60x objective described above.

### A. calliptera and R. sp. chillingali staining and imaging

Cichlid fish adults were kept at 26ºC, under a 12:12 hour day-night cycle. Embryos were fixed overnight, at 4ºC, in 4% formaldehyde. Following fixation, embryos were rinsed in PBST (1x PBS, Sigma Aldrich, and 0.1% Tween-20) three times before being removed from the yolk prior to staining. Embryos were permeabilised for two days total in PBS + 1% Triton-X100, washing the embryos four times total. Embryos were then stained overnight in 100*µ*L PBST, 10*µ*L Phalloidin-CF568 (Biotin, BT00044-T), and 0.5*µ*L DAPI (Biotin, BT40011), before being rinsed 3x in PBST, and mounted in 90% glycerol. *A. calliptera* were imaged using an LSM880 under a 40x water immersion lens (C-Apochromat 40x/1.2 W Korr FCS M27) and *R. sp. chillingali* were imaged on an Olympus FV3000 under a 30x silicon oil lens (UPlanSApo 30x silicon oil, 1.05NA).

### Active trapping

Active trapping of embryos within the chorion was carried out by immersing 24 hpf embryos in 0.006% bleach (30*µ*L 10% hypochlorite per 50mls of E3 media) for 5 min. Embryos were subsequently washed three times in fresh E3 which was changed daily. Full protocols can be found in Ref (99).

### Hardware

For image processing, data analysis and visualization three computing systems were used depending on the computational demand required. A workstation Intel Xeon, 512 GB RAM, Nvidia Quadro P2000 GPU 24 GB, running Windows 10; a workstation Intel Xeon, 62 GB RAM, Nvidia Quadro P6000 GPU 48 GB, Ubuntu 20.04 and a MacBook Pro, M2 Pro apple silicon, 32 GB RAM.

## Acknowledgments

This work was funded by BBRSC Responsive Mode Grant BB/W006944/1 and UKRI Physics of Life EP/W023865/1. We thank the Warwick imaging facility (CAMDU) and Warwick BSU for support. RGP was supported by an Australian Research Council (ARC) Laureate Fellowship (FL210100107) and TEH by the National Health and Medical Research Council of Australia (Ideas Grant 2027559). We thank Jianmin Yin for providing the data on *fused somite* and Sunandan Dhar for the actin imaging. This research was supported in part by NSF grants PHY-2309135 and PHY-2309135 and the Gordon and Betty Moore Foundation Grant No. 2919.02 to the Kavli Institute for Theoretical Physics (KITP), where S.T. and T.E.S. developed part of this work.

