## Supplementary Information for "A structural transition ensures robust formation of skeletal muscle"

DRAFT

### Contents


- Supplementary Figures
- Supplementary Movie Legends

### Supplementary Figures

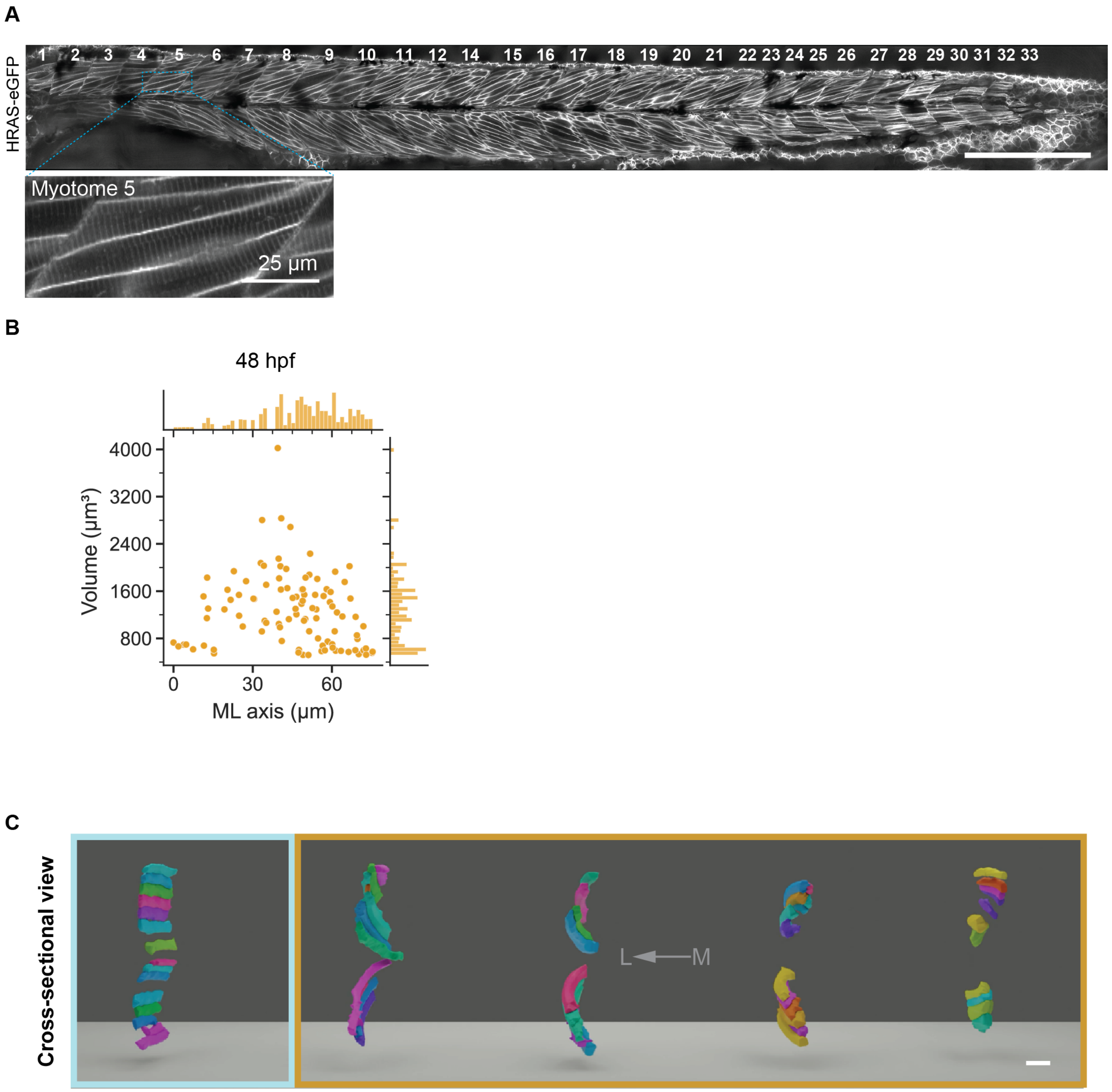


**Supporting Figure 1.** **A)** Flattened 48 hpf embryo showing somite segments from 1 (oldest, most anterior) to 33 (youngest, most posterior). Inset shows sarcomere formation in more mature myotome segments. Scale bar $200\mu m$. **B)** Quantification of cell volume in developing myotome as function of ML position in wt embryos at 48 hpf. n = 100 cells. **C)** Cross-sectional view of cell segmentation along ML-axis of different layers at 48 hpf. Scale bar = 25 $\mu m$.

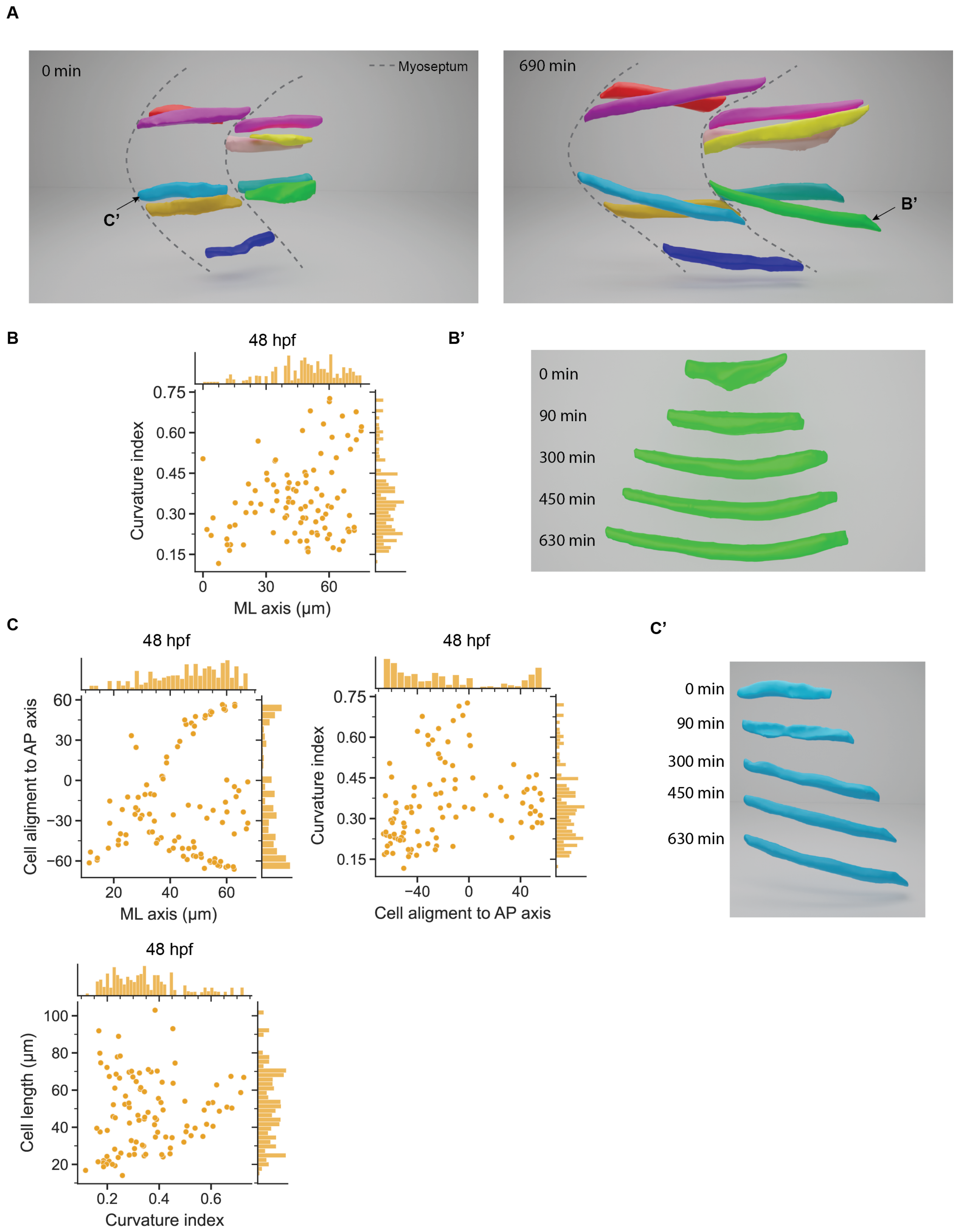


**Supporting Figure 2.** **A)** Comparison of cell orientation at beginning of elongation (left) and after the initial wave of fibre morphogenesis is complete (right). Cells initially align with the AP axis. **B)** Left: quantification of cell curvature at 48 hpf with respect to ML position. Right: segmentation of example cell highlighting how curvature emerges during fibre elongation. n = 100 cells. **C)** Left: Cell alignment with respect to ML axis at 48 hpf. Centre: Curvature with respect to AP axis at 48 hpf. Right: Example cell segmentation highlighting onset of skew during fibre elongation. Bottom: Cell length with respect to the cell curvature index. n = 100 cells.

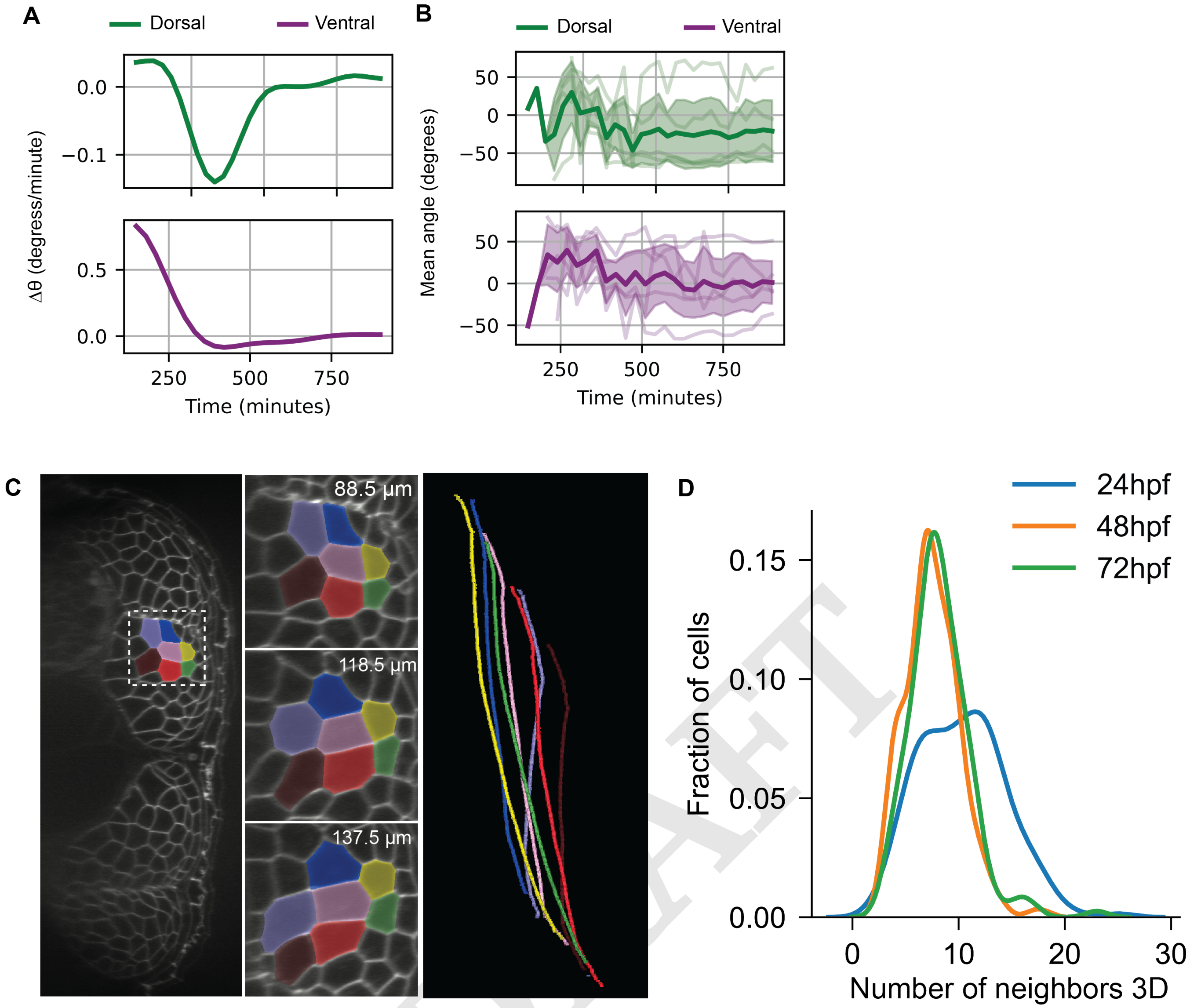


**Supporting Figure 3.** **A)** Quantification of cell twist onset in cells positioned dorsal (top) and ventral (bottom) in the developing myotome. $n = 5$ cells per zone from 2 myotomes. **B)** Average behaviour of twist over time. $n = 5$ cells per zone from 2 myotomes. **C)** Example of group of cells undergoing twist at 48phf. Right: centroid position of cells along AP-axis. **D)** Neighbour number distribution of future muscle fibres at 24 ($n = 329$ cells), 48 ($n = 341$ cells) and 72 hpf ($n = 328$ cells). $N = 3$ embryos per condition).

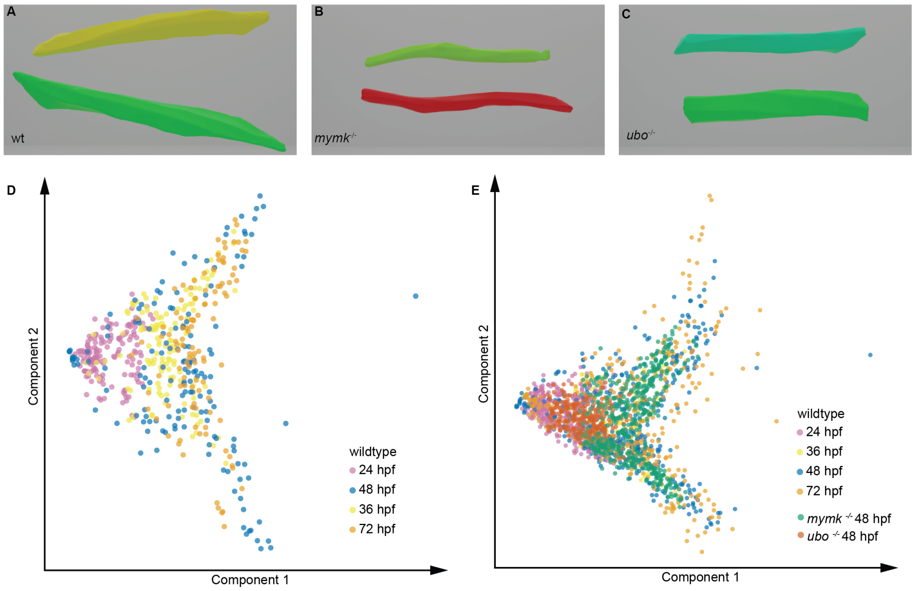


**Supporting Figure 4.** **A)** Example cell segmentations from dorsal and ventral domains of 48 hpf myotome segment. **B)** Example cell segmentations from similar location as (A) in a *mymk* mutant embryo. **C)** Example cell segmentations from similar location as (A) in a *ubo* mutant embryo. **D-E)** Data from Figure 4, Figure 5E plotted using their first 2 PCA components.

DRAFT

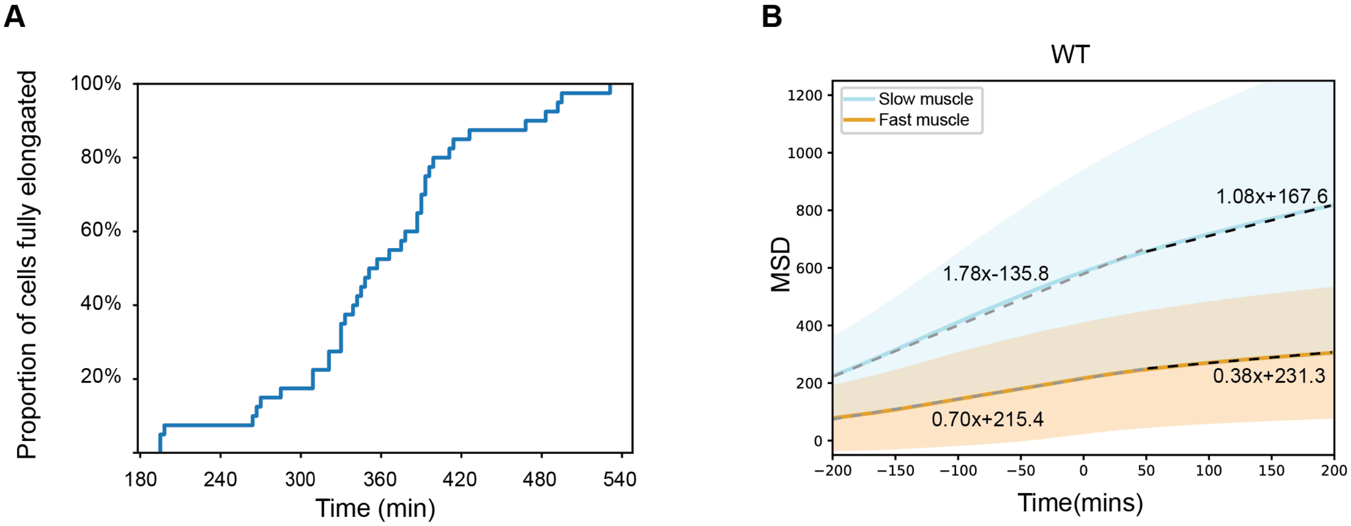


**Supporting Figure 5.** **A)** Proportion of future fast fibre myocytes that have extended fully between the anterior and posterior boundaries of the myotome segment. 40 cells from 2 myotomes. Time 0 min corresponds to somite segmentation. **B)** MSD as Fig. 6A but with time defined as $t = 0$ for each cell when fusion occurs. Equations represent best linear fits to data before and after $t = 0$ for fast and slow fibres.

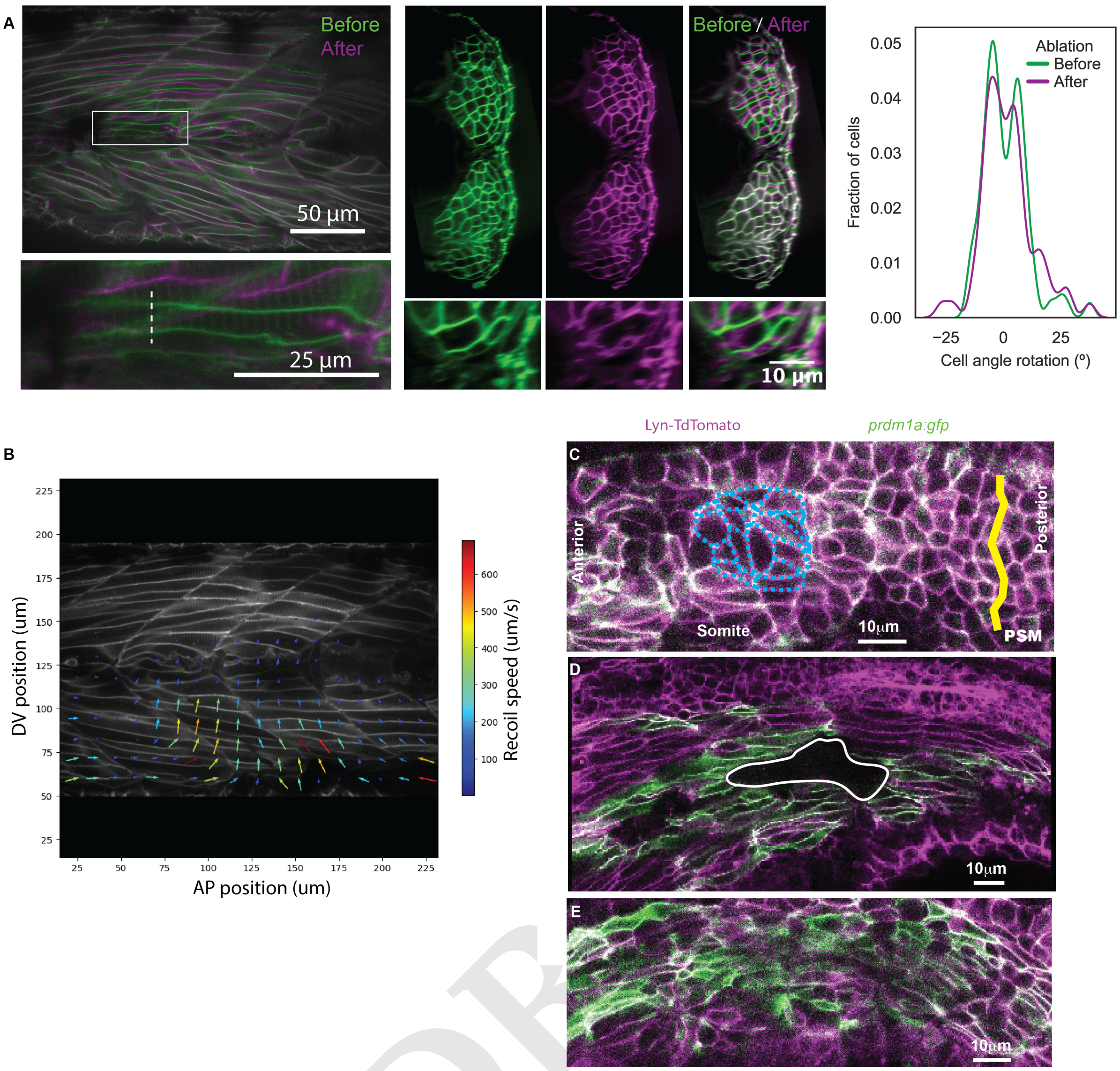


**Supporting Figure 6.** **A)** Comparison of muscle structure before and after ablation of myoseptum. Right: quantification of change in cell rotation angle. Before: $n = 76$ cells; after: $n = 66$ cells. **B)** Flow fields after ablation of a future fast muscle fibre. Only nearby cells respond, with cells across the myoseptum showing little response. **C)** *fused somite* mutant embryo imaged at 18-22 somite stage. The putative PSM boundary highlighted by yellow line. Example cell outlines highlighted by blue dashed lines. There is no clear cell orientation with AP axis. **D)** Example of nonconfluent somite region (white outline) from a *fused somite* mutant embryo. **E** Slow muscles, marked by SMHC::GFP do not form a clear continuous layer with *fused somite* mutant embryos.

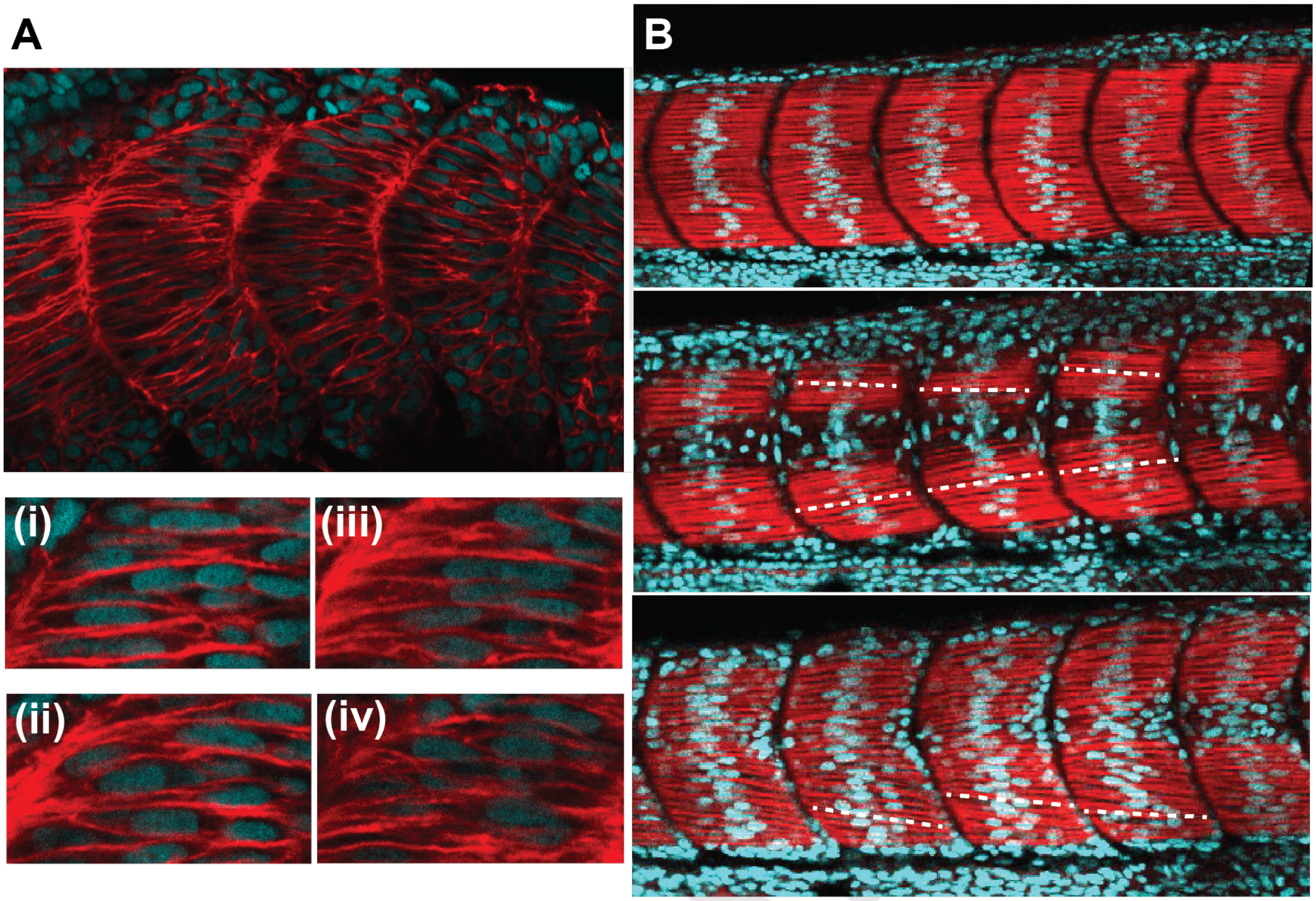


**Supporting Figure 7.** Supporting Discussion. **A)** *A. calliptera* embryo at stage 13 stained with DAPI (cyan) and phalloidin (red). (i)-(iv) show the same section of muscle at different ML positions. The twisting of the cell membrane is apparent. **B)** *R. chillingal* embryo at stage 14 stained with DAPI (cyan) and phalloidin (red). Top: mid ML-plane. Centre: near medial surface. Bottom: near lateral surface. Cell alignment with AP axis demonstrated with dashed white lines. Embryo staging based on Marconi *et al.*, Evolution and Development 2023.

DRAFT

### Supplementary Movie Legends

**Movie S1**: Wildtype embryo expressing Lyn-GFP at somite stage 21.

**Movie S2**: Segmentation of wildtype myotome segment at 48 hpf, visualised in Blender.

**Movie S3**: Visualisation of twisted cell at 48 hpf.

**Movie S4**: Time course of selected segmented cells from 24 to 48 hpf.

**Movie S5**: Demonstration of software for interrogating the spherical harmonic representation of wildtype cells at different developmental time periods. Left shows the UMAP reduction of the spherical harmonic representation of the cell shapes and right the midplane of the selected segments.

**Movie S6**: Time course of cell shape through the UMAP reduction of the spherical harmonic representation of each cell.

**Movie S7**: Cell tracks of wildtype (left) and $mymk^{MO}$ treated embryos (right) starting from somite generation.

**Movie S8**: Ablation of muscle pioneers before myocyte elongation.

**Movie S9**: Ablation of muscle pioneers after myotube formation.

**Movie S10**: Ablation muscle progenitors after myotube formation.

DRAFT
